# The Genome Sequence Archive Family: Towards Explosive Data Growth and Diverse Data Types

**DOI:** 10.1101/2021.06.29.449849

**Authors:** Tingting Chen, Xu Chen, Sisi Zhang, Junwei Zhu, Bixia Tang, Anke Wang, Lili Dong, Zhewen Zhang, Caixia Yu, Yanling Sun, Lianjiang Chi, Huanxin Chen, Shuang Zhai, Yubin Sun, Li Lan, Xin Zhang, Jingfa Xiao, Yiming Bao, Yanqing Wang, Zhang Zhang, Wenming Zhao

**Author notes:** Corresponding author(s). (Zhao W), (Zhang Z), (Wang Y). Equal contribution (Tingting Chen, Xu Chen, Sisi Zhang, Junwei Zhu).

## Abstract

The Genome Sequence Archive (GSA) is a data repository for archiving raw sequence data, which provides data storing and sharing services for worldwide scientific communities. Considering explosive data growth with diverse data types, here we present the GSA family by expanding into a set of resources for raw data archive with different purposes, namely, GSA (https://ngdc.cncb.ac.cn/gsa/), GSA for Human (GSA-Human, https://ngdc.cncb.ac.cn/gsa-human/), and Open Archive for Miscellaneous Data (OMIX, https://ngdc.cncb.ac.cn/omix/). Compared with the 2017 version, GSA has been significantly updated in data model, online functionalities, and web interfaces. GSA-Human, as a new partner of GSA, is a data repository specialized in human genetics-related data with controlled access and security. OMIX, as a critical complement to the two resources mentioned above, is an open archive for miscellaneous data. Together, all these resources form a family of resources dedicated to archiving explosive data with diverse types, accept data submissions from all over the world and provide free open access to all publicly available data in support of worldwide research activities.

## Introduction

The Genome Sequence Archive [1] (GSA, https://ngdc.cncb.ac.cn/gsa) is a public archive of raw sequence data in the National Genomics Data Center (NGDC) [2–4], part of the China National Center for Bioinformation (CNCB). GSA accepts worldwide data submissions, performs data curation and quality control for all submitted data, and provides free open access to all publicly available data without unnecessary restrictions. Since its inception in 2015, GSA has been broadly supported and endorsed by the scientific community, as testified by a total of 324,325 experiments, 371,973 runs and 8526 TB files submitted by 1530 users from 385 institutions and reported in 634 research articles and 239 scientific journals (as of June 2021). Importantly, GSA serves as one of the core resources in CNCB-NGDC that has stable state funding in biological data management, thus ensuring long-term persistence and preservation of submitted datasets.

Due to the rapid development of sequencing technologies towards higher throughput and lower cost as well as their wider applications in biomedical research, a large number of multi-omics data have been produced at ever-increasing rates and scales, provoking two major challenges for raw data management in GSA. For one thing, several large-scale sequencing projects (such as Earth BioGenome Project [5], Dog 10K Project [6], Protist 10000 Genomes Project [7]) have been carried out over the past several years, leading to different types of raw sequence data generated around the global and accordingly requiring a suite of web services for massive data submission and deposition. For another, studies on human population genomics and precision medicine have produced millions of personal genome sequences associated with clinical information, requiring controlled access and security management, which is critically vital in promoting human healthcare and precise medical treatment and advancing big-data-driven scientific research, while protecting data privacy. These challenges are particularly crucial in China since it not only features the largest population in the world and rich biodiversity resources, but also has a formidable capacity in genome sequencing throughout the country.

To address these challenges, here we provide a family of resources for raw data archive and management, including an updated version of GSA and two newly developed partner resources, namely, GSA for Human (GSA-Human, https://ngdc.cncb.ac.cn/gsa-human) and Open Archive for Miscellaneous Data (OMIX, https://ngdc.cncb.ac.cn/omix). Specially, we updated GSA with significant improvements on data model, online functionalities and web interfaces. As an important partner to GSA that provides open access to all released data, GSA-Human features controlled-access and security services for human genetics-related data, which is compatible well with the database of Genotypes and Phenotypes (dbGaP) [8] in the National Center for Biotechnology Information (NCBI) [9] and the European Genome-phenome Archive (EGA) [10] in the European Bioinformatics Institute (EBI) [11]. But GSA-human is different from dbGaP and EGA; the former is mainly used to archive and store raw sequence data, while the latter not only archive raw sequence data, but also archive phenotypic data. In addition, OMIX (https://ngdc.cncb.ac.cn/omix/), as a critical complement to the above two resources, is an open archive for miscellaneous data that are unsuitable to store in GSA, GSA-Human or other databases at CNCB-NGDC. Together, all these resources form a family of resources dedicated to archiving explosive data with diverse types.

## Archival resources

GSA, built based on the INSDC (International Nucleotide Sequence Database Collaboration) [12] data standards and structures, is a public data repository for archiving raw sequence reads. Over the past several years, GSA has been frequently and considerably updated since its establishment in 2015, with significant improvements in data structure, online submission, quality control, and web functionalities (**Table 1**). First, data structure has been significantly changed (Figure 1); BioProject (https://ngdc.cncb.ac.cn/bioproject/) and BioSample (https://ngdc.cncb.ac.cn/biosample/) have been separated from GSA, serving as independent meta-information databases and acting as an organizational framework to provide centralized access to descriptive metadata about research projects and samples, respectively. Second, to help users submit massive data with different types, more sequencing platforms, sample types, and file formats were acceptable, and importantly, batch submission of multiple experiments and runs was enabled in the updated version of GSA. In addition, to provide users with convenient services for uploading raw sequence files, GSA not only provides an FTP server but also equips with Aspera (https://www.ibm.com/products/aspera) to realize high-speed data transmission. Third, GSA was greatly enhanced by improving the expert curation process and integrating an automated quality control pipeline, with the aim to provide value-added services for archiving high-quality data. Fourth, multiple web functionalities for bilingual support (both English and Chinese), online documentation, data statistics and visualization charts, were updated/newly added. Taken together, the updated version of GSA is more efficient and friendly in big omics-data submission, deposition and management.

**Table 1.**
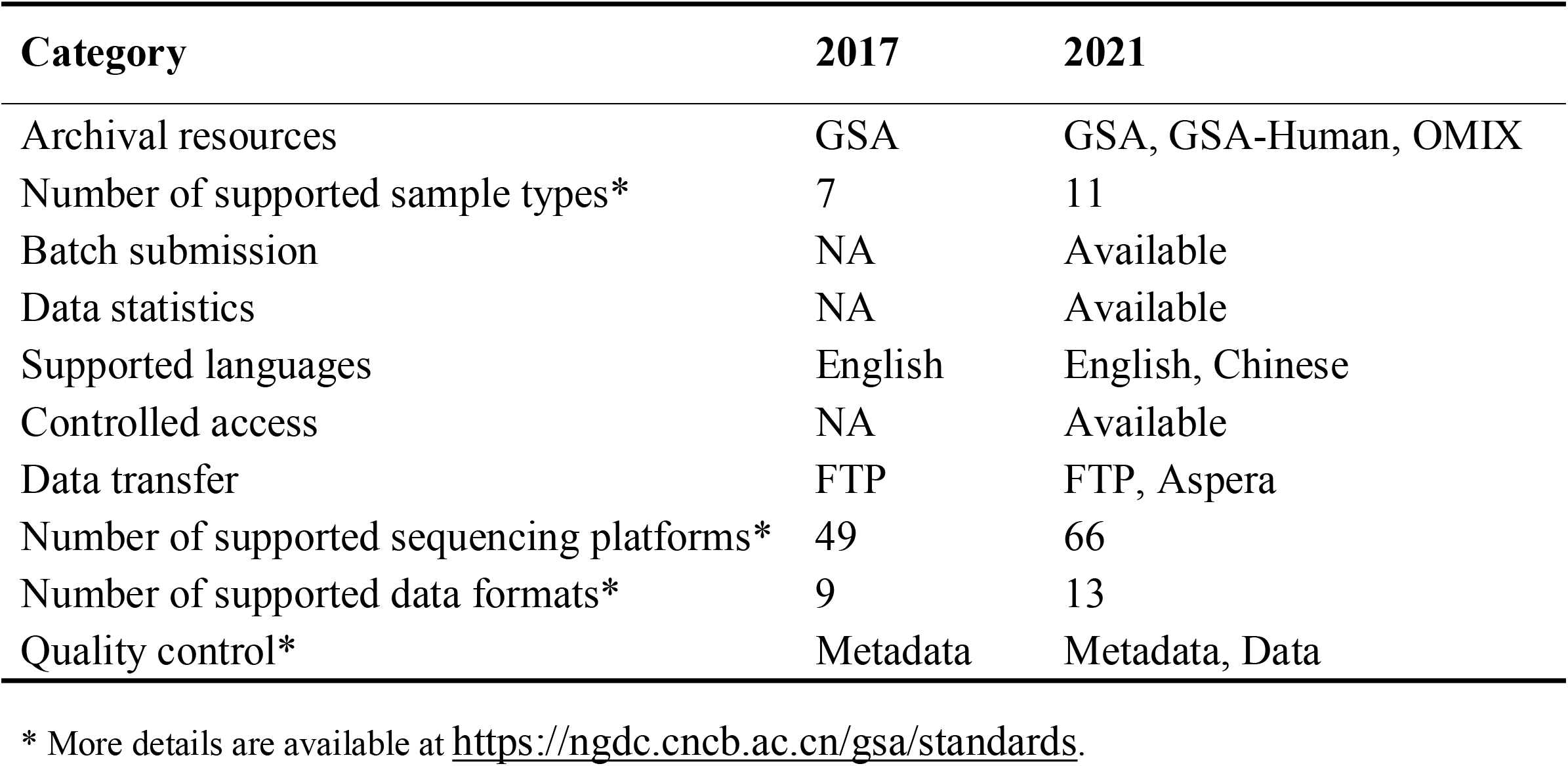
Comparison between GSA in 2017 and the GSA family in 2021.

| **Category** | **2017** | **2021** |
| --- | --- | --- |
| Archival resources | GSA | GSA, GSA-Human, OMIX |
| Number of supported sample types* | 7 | 11 |
| Batch submission | NA | Available |
| Data statistics | NA | Available |
| Supported languages | English | English, Chinese |
| Controlled access | NA | Available |
| Data transfer | FTP | FTP, Aspera |
| Number of supported sequencing platforms* | 49 | 66 |
| Number of supported data formats* | 9 | 13 |
| Quality control* | Metadata | Metadata, Data |

\* More details are available at https://ngdc.cncb.ac.cn/gsa/standards.

**Figure 1.**
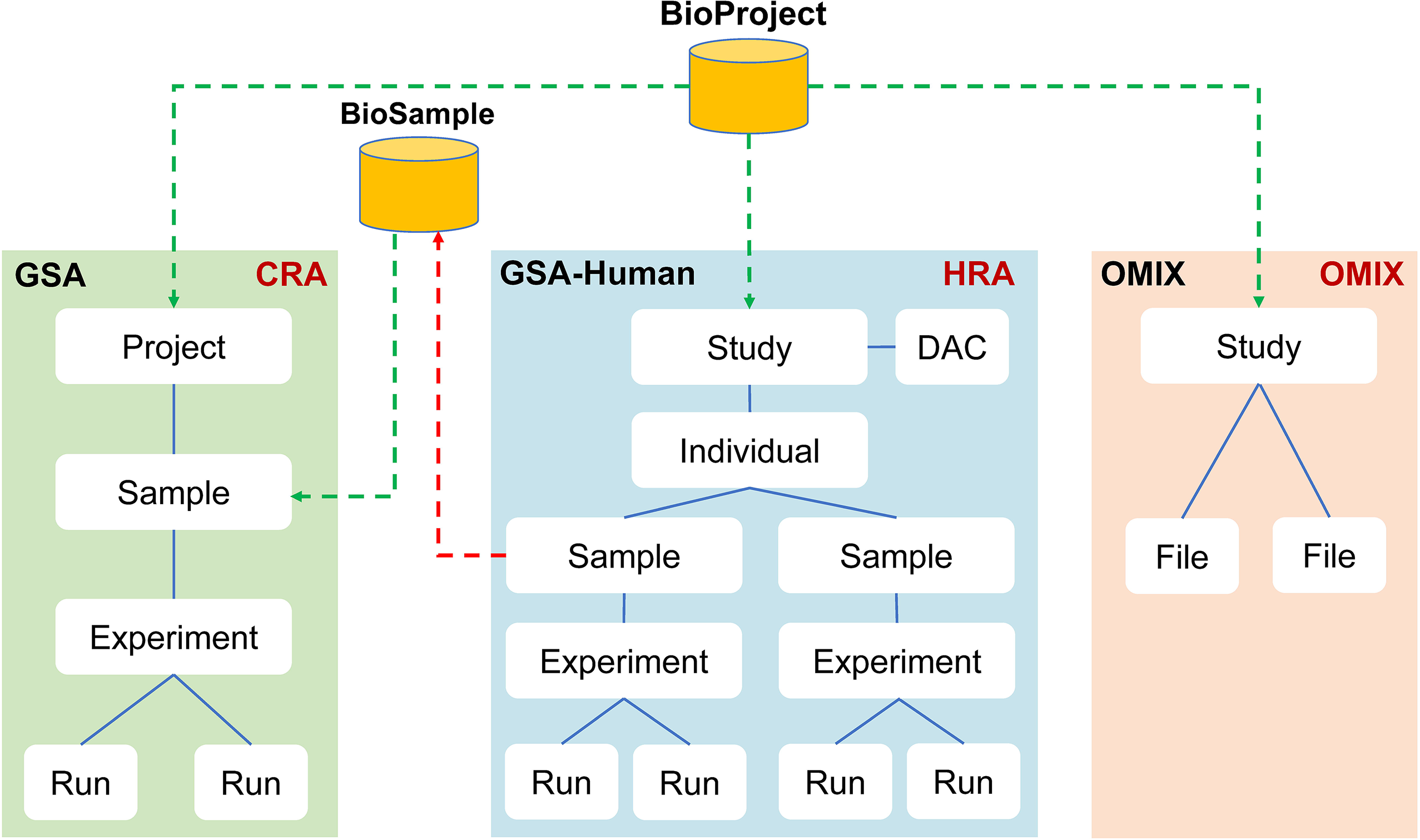
Data model of the GSA family. GSA data structure has been significantly changed. BioProject and BioSample have been separated from GSA, serving as independent meta-information databases and acting as an organizational framework to provide centralized access to descriptive metadata about research projects and samples, respectively. GSA-Human is used to archive human genetic resources data and OMIX is used for various non-sequencing types of data management.

GSA-Human, established in April 2018, is a data repository specialized in the secure management of human genetics-related data. It accepts submissions of various studies, including disease, cohort, cell line, clinical pathogen and human associated metagenome. GSA-Human uses the “individual” to organize its metadata and sequence reads and provides two different data access mechanisms: open access and controlled-access. Open access means that all data are public for global researchers, whereas controlled-access means that data can be downloadable only after being authorized by the Data Access Committee (DAC) that is responsible for authorizing/declining data access to data requester. Therefore, GSA-Human provides a series of data services including access control, data request, access authorization/decline, and security management.

OMIX, as a new member of the archival resources in CNCB-NGDC, aims to meet users’ needs for submitting various types of data other than sequences. It collects not only raw data from transcriptome, epigenome, and microarray, but also functional data such as lipidome, metabolome, proteome, and even data like clinical information, demographic data, questionnaire and so on. With the concise interface and simplified submission process, OMIX enables data submission and deposition very easy. Of note, similar to GSA-Human, OMIX has a data security management strategy for human genetic data. Any controlled-access dataset in OMIX can be accessed only with the permission of the original data submitter/owner.

### Data submission and retrieval

Data submission to the GSA family is aided by a series of web services, including BIG Single Sign-On (SSO; https://ngdc.cncb.ac.cn/sso/) that is a user access control system and BIG Submission portal (BIG Sub; https://ngdc.cncb.ac.cn/gsub/) that is a unified one-stop portal providing submission services for a variety of database resources in CNCB-NGDC. To submit data to the GSA family, user needs to register an account and log into any database via SSO that can help user gain access to multiple independent systems with a single ID and password.

Overall, the GSA family provides a suite of services for data retrieval, download and access. Public data in these resources can be retrieved via BIG Search (https://ngdc.cncb.ac.cn/search/), a scalable text search engine that performs more powerful data retrieval and analytical capabilities. All released data are publicly accessible and downloadable via FTP and HTTP, but controlled data in GSA-Human and OMIX require access permission. To access the controlled data, requester needs to create a request and send required documents for data access. Once the request has been reviewed and approved, the requester gains the access to the data.

### Data statistics

The GSA family has received a large number of data submissions with explosive growth in data and users, thus exhibiting their important roles in raw data management (**Figure 1** and **Table 2**). The volume of archived data has increased by more than 40 times, compared to the 200 TB archived in the previous release of GSA [1]. Till June 2021, GSA and GSA-Human have collected 324,325 Experiments, 371,973 Runs and more than 8.5 PB of data submitted from 1530 submitters of 385 organizations (Figure 2A). In particular, GSA-Human has archived 61,225 individuals and housed 4.9 PB of raw sequence data within less than one year, clearly showing that human genetic data are growing at an unprecedented rate and scale. More importantly, GSA-Human has received a total of 721 access requests from 485 requesters, with 178 requests approved till June 2021. Regarding the trend of archived data over time, it is observed that it took about three years to accumulate the first PB of data and currently reaches to 8.5 PB in just over two and a half years, with a formidably dramatic decrease in days for data accumulation (Figure 2B). Strikingly, the third PB volume took only 30 days, principally contributed by a large-scale sequencing project [13] with 344 TB of data archived. Meanwhile, the number of species involved is also on a rapid increase, from 80 in December 2016 to more than 1000 at present. Also, albeit newly established, OMIX has collected 160 files of 801 GB.

**Table 2.** Data items of the GSA family.

| **Item\*** | **GSA** | **GSA-Human** | **OMIX** | **Total** |
| --- | --- | --- | --- | --- |
| BioProjects | 2398 | 537 | 83 | 2920 |
| Individuals | / | 61,225 | / | 61,225 |
| BioSamples | 241,360 | 125,715 | / | 367,075 |
| Experiments | 178,670 | 145,655 | / | 324,325 |
| Runs | 195,298 | 176,675 | / | 371,973 |
| File size (Tbyte) | 3545 | 4980 | 0.888 | 8526 |
| Registered users |  | 4610 |  |  |

\* All statistics were derived from the GSA family as of June 2021.

**Figure 2.**
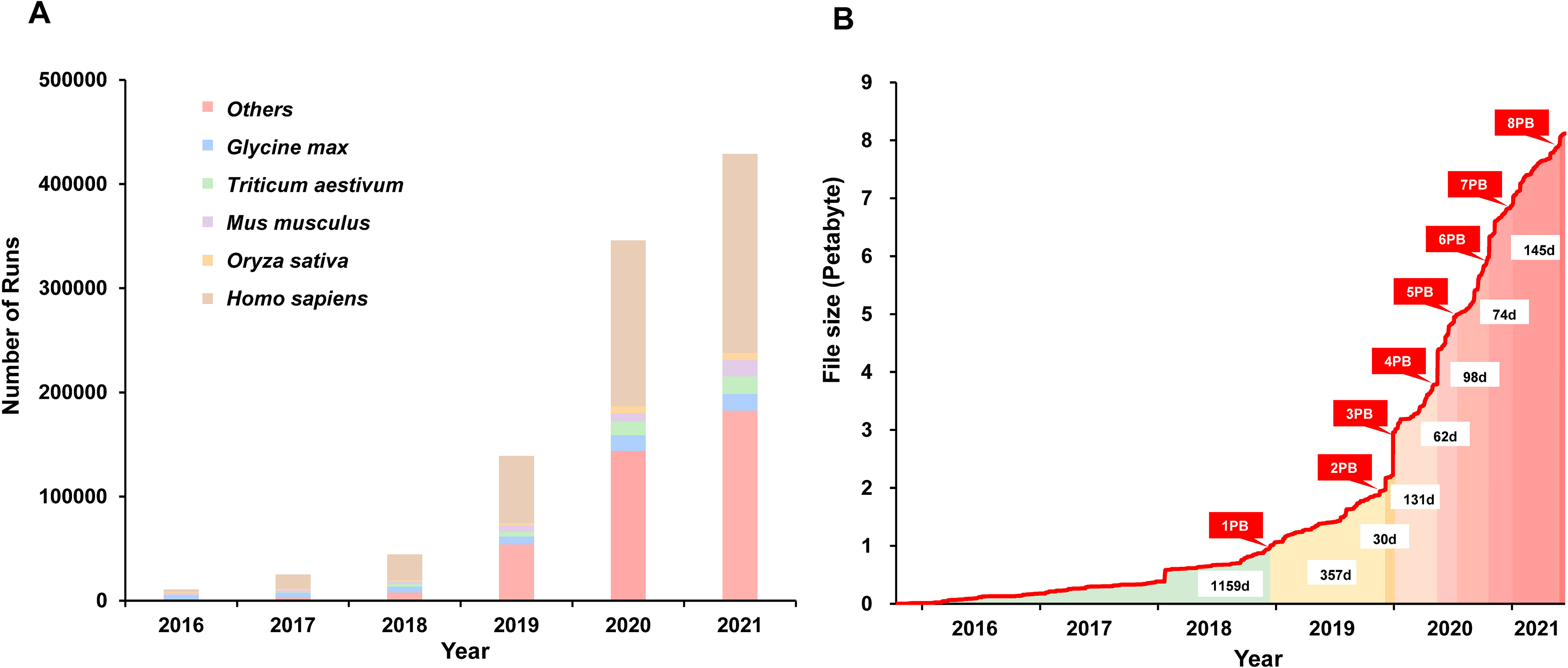
Data statistics of the GSA family. Number of runs accumulated from 2016 to 2021, with five major species indicated. Trend of submitted data volume in association with days involved. All statistics were derived from GSA and GSA-Human as of June 2021.

Currently, the GSA family has more than 5377 registered users and has been visited by 648,274 unique IPs from 111 countries/regions, with a total of 35,010,529 page views and an average of 4 TB of downloads per day. Data housed in these resources have been reported in more than 239 scientific journals(https://ngdc.cncb.ac.cn/gsa/statistics?active=journals), including Cell, Genome Research, Genomics Proteomics Bioinformatics, Nature, Plant Cell and PNAS. More importantly, with frequent updates and improvements in the past several years, GSA has been recognized as one of the certified repositories in FAIRsharing.org and re3data.org, and therefore meets the requirement as a supported repository by Elsevier, Taylor & Francis, and Wiley. More detailed statistics can be found online at https://ngdc.cncb.ac.cn/gsa/standards.

### Future directions

The explosive volume of raw data submitted to the GSA family is still on the increase, posing significant challenges to handle and share such big data [14]. Nowadays, CNCB-NGDC, hosting a suite of database resources including the GSA family, is going to be enhanced by national big data infrastructure, with stable governmental funding investment in upgrading storage, computing and network resources, thus providing fundamental support in raw data archive and management of the GSA family. In addition, our future efforts will be made in continuous optimization of data models and curation processes in evolution of users’ needs, establishment of cloud infrastructure for big data storage, and development of a variety of tools to facilitate big data submission and high-speed transfer. To make effective use of human genetic data and promote precision healthcare and treatment, efforts will also be devoted to optimizing procedures and mechanisms to enable data sharing with controlled access and security by conforming to applicable regulations and ethical norms. We also advocate worldwide collaborations in developing data standards, tools and approaches towards global biodiversity & health big data sharing (BHBD alliance; http://bhbd-alliance.org/).

### CRediT author statement

Tingting Chen: Investigation, Methodology, Data Curation, Writing - Original Draft. Xu Chen: Software. Sisi Zhang: Investigation, Methodology, Data Curation, Writing - Original Draft. Junwei Zhu: Software. Bixia Tang: Software. Anke Wang: Writing - Original Draft, Software. Lili Dong: Data Curation. Zhewen Zhang: Data Curation. Caixia Yu: Data Curation. Yanling Sun: Data Curation. Lianjiang Chi: Software. Huanxin Chen: Resources. Shuang Zhai: Resources. Yubin Sun: Resources. Li Lan: Resources. Xin Zhang: Resources. Jingfa Xiao: Writing - Review & Editing. Yiming Bao: Conceptualization, Writing - Review & Editing, Funding acquisition. Yanqing Wang: Conceptualization, Investigation, Methodology, Software, Writing - Review & Editing, Project administration. Zhang Zhang: Conceptualization, Writing - Review & Editing, Funding acquisition. Wenming Zhao: Conceptualization, Methodology, Writing - Review & Editing, Supervision, Funding acquisition.

## Acknowledgments

We sincerely thank Prof. Jingchu Luo and Prof. Weimin Zhu for their valuable suggestions and a number of users for their contributions to the data submission. We also thank Changrui Feng and Zhuojing Fan for their assistance in drawing the figures of the manuscript. This work was supported by grants from Strategic Priority Research Program of Chinese Academy of Sciences [XDB38060100 and XDB38030200 to Y.B.; XDB38050300 to W.Z.; XDB38030400 to J.X.; XDA19050302 to Z.Z.]; National Key Research and Development Program of China [2016YFC0901603 to W.Z.; 2017YFC1201202 to Y.W.; 2020YFC0847000 and 2018YFD1000505 to W.Z.; 2016YFE0206600 to Y.B.; 2017YFC0907502 to Z.Z.]; The 13th Five-year Informatization Plan of Chinese Academy of Sciences [XXH13505-05 to Y.B.]; Genomics Data Center Construction of Chinese Academy of Sciences [XXH-13514-0202 to Y.B.]; Open Biodiversity and Health Big Data Programme of IUBS [to Y.B.]; The Professional Association of the Alliance of International Science Organizations [ANSO-PA-2020-07 to Y.B.]; National Natural Science Foundation of China [32030021 and 31871328 to Z.Z.]; International Partnership Program of the Chinese Academy of Sciences [153F11KYSB20160008 to Z.Z.].

